# Daily Activity Profiles over the Lifespan of Female Medflies as Biomarkers of Aging and Longevity

**DOI:** 10.1101/2023.06.07.543948

**Authors:** Han Chen, Hans-Georg Müller, Vasilis G. Rodovitis, Nikos T. Papadopoulos, James R. Carey

## Abstract

The relationship between the early age activity of Mediterranean fruit flies or other fruit flies and their lifespan has not been much studied, in contrast to the connections between lifespan and diet, sexual signaling and reproduction. The objective of this study is to assess intraday and day-to-day activity profiles of female Mediterranean fruit flies and their role as biomarker of longevity as well as to explore the relationships between these activity profiles, diet and age-at-death throughout the lifespan. Three distinct patterns of activity variations in early age activity profiles can be distinguished. A low-caloric diet is associated with a delayed activity peak, while a high-caloric diet is linked with an earlier activity peak. We find that age-at-death of individual medflies is connected to their activity profiles in early life. An increased risk of mortality is associated with increased activity in early age, as well as with a higher contrast between daytime and nighttime activity. Conversely, medflies are more likely to have a longer lifespan when they are fed a medium caloric diet and when their daily activity is more evenly distributed across the early age span and between daytime and nighttime. The before-death activity profile of medflies displays two characteristic before-death patterns, where one pattern is characterized by slowly declining daily activity and the other by a sudden decline in activity that is followed by death.

## 1 Introduction

The study of the mortality and longevity of fruit flies is well established in the biodemographic literature. For instance, Zhang et al. (2006) provided evidence that high levels of recent calling activity in male medflies are linked to longer remaining lifespans. Müller et al. (2001) demonstrated that the rate of exponential decrease in egg laying by female medflies is predictive for their remaining lifespans while Meng et al. (2021) showed that egg-laying patterns of fruit flies in terminal segments could be distinguished from patterns in non-terminal segments. Harshman and Zera (2007) investigated the connection between early-life increments in reproduction and later survival, and explored the biological mechanisms behind this relationship, while Harwood et al. (2013) examined the impact of diet and host access on the lifespan of the fruit fly.

The literature on fly activity tracking is extensive. For example, Shaw et al. (2019) analyzed the daily activity patterns of Drosophila suzukii, considering factors such as gender, physical conditions, and the circadian clock on these patterns. Kaladchibachi et al. (2019) analyzed the circadian patterns of activity change in female Drosophila ananassae and studied its relationship with aging. Although previous studies aimed at investigating this relationship, they have been limited in their methodology and did not feature continuous comprehensive monitoring (Sohal and Buchan 1981; Lints et al. 1984; Sohal 1985; Le Bourg 1987). For instance, Sohal and Buchan (1981) stated that the lifespan of individual flies corresponds to their levels of physical activity observed between day 4 and day 7, while Le Bourg (1987) found no significant correlation between lifespan and average locomotor activity observed once per week. However, these studies used an aggregated activity score as a measure of activity, which does not provide a comprehensive quantification of the daily activity profile. Furthermore, none of these studies monitored activity longitudinally with a 24/7 activity monitor throughout the lifetime of each individual, which provides a more accurate and detailed representation of activity and thus the relationship between movement activity levels and lifespan. Additionally, the correlation calculation used in these studies was not amenable for the exploration of more complex dependencies between lifespan and activity levels.

In our study, we used the LAM25 system to obtain repeated observations of the 24-hour movement activity of Mediterranean fruit flies. The system records the movement of each fly as it crosses a barrier. We model the intraday and day-to-day activity profiles and their relationship with the remaining lifespan of each individual through functional data analysis for repeated observations. This approach models the activity profiles as manifestations of an underlying function-valued stochastic process (Wang et al. 2016; Chen and Müller 2012; Chen et al. 2017). In our analysis, the longitudinal time is the age of the fly (in days) and the repeated observations consist of the daily activity records that are recorded in terms of within day time (0-24 hours).

The goals of our investigation are (1) To study the relationship between activity and aging dynamics and the role of activity as a biomarker of longevity, for which we utilize function-valued principal component analysis (Chen and Müller (2012)) for repeated functional data to construct the principal components of the underlying activity process, separating the intra-day and inter-day components. (2) To investigate the connection between early-age activity and lifespan at the subject level. For this we use functional linear regression to predict an individual’s remaining lifespan, given the activity profile and diet. (3) To explore before-death pattersn of decline in activity.

Three distinct patterns of variation in the early-age activity profiles are identified in the study, and their connection to diet is investigated. Results show that a medium nutrition level contributes to a relatively evenly distributed activity profile, while a low nutrition level is associated with a relatively late activity peak and a high nutrition level with a relatively early activity peak. In terms of the connection between early age activity profile and lifespan, we find that increased mortality risk is linked to higher activity levels and a larger contrast between daytime and nighttime activity, particularly in individuals with a high nutrition level. Cluster analysis identified two patterns of before-death activity profiles, one corresponding to continuous dwindling of activity before death and another one characterized by a rapid decline of activity in a short time period.

## 2 Data Description

Three locomotory activity Monitor-LAM25 systems were used to record the locomotory activity patterns of the adult medflies. In this system, each fly was put in its own glass tube (25 mm diameter, 125 mm length), where three ray rings of infrared light monitors at three different planes were placed (close to the two ends of the tube and one in the middle). The activity of each tube was measured every minute, as the number of times the fly passed through an infrared beam. The agar-based gel diet consisting of sugar, yeast, hydrolysate, agar, nipagin and water (4:1:0.2:0.1:20) was prepared to provide adults with both nutrients and water simultaneously. We used three different gel diets which differ in the sugar and yeast hydrolysate content in the gel (50%, 20% and 10%, represented by treatment C-50, C-20 and C-10 accordingly). The gel diet was replaced every four days to avoid dehydration. Females’ mortality was recorded daily with visual observations until the death of all individuals.

In our analysis, individual activities from the record of the middle monitor were used and aggregated per hour for the total number of 96 tubes under three different treatment levels (C-10, C-20, C-50). Each treatment level corresponds to 32 tubes. See Figure 7, 8 and 9 for the daily activities under each treatment together with the individual lifespan.

## 3 Statistical Methods

### 3.1 Function-valued Stochastic Process

Denote the time coordinate of the calendar days by *t* ∈ 𝒯 and the repeated measurement coordinate of hours within a day by *s* ∈ 𝒮. Let *X*(*s, t*) to be the function-valued stochastic process representing the activity level at day *t* and hour *s*. Mean function and the covariance function of the process can be defined as

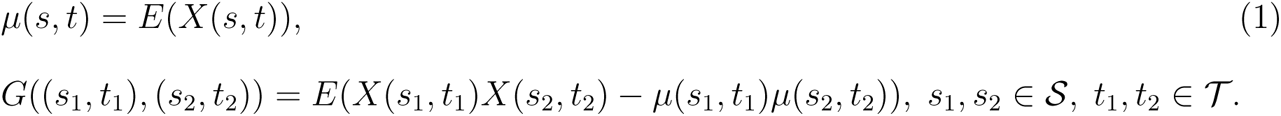

Mercer’s theorem implies the spectral decomposition of the covariance surface *G* that

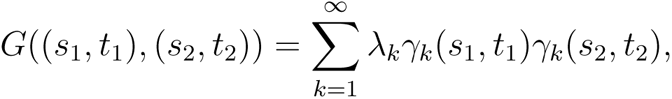

where *λ_k_* and *γ_k_* are eigenvalues and two-dimensional eigenfunctions of the covariance operator with covariance kernel *G*. By the Karhunen-Lòeve expansion (Karhunen 1946), one can represent *X*(*s, t*) through Two-dimensional Functional Principal Component Analysis (FPCA) as

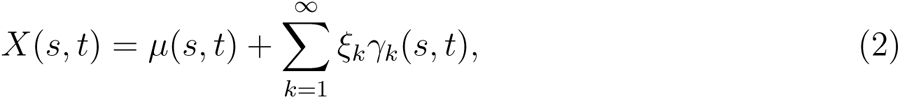

where 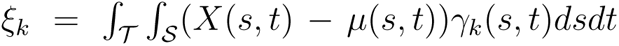 are principal components of the process. These components are zero mean uncorrelated random variables representing the fluctuations of the process *X*(*s, t*) around the mean function *µ*(*s, t*). In practice, one works with the first *K* components as the approximated process that 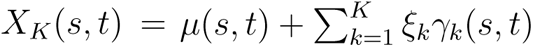. We choose *K* = 3 in the following analysis.

Modes of variation (Castro et al. 1986) are a common tool to interpret and visualize the FPCA. They focus on the contributions of each eigenfunction to the stochastic behavior of the underlying process. The *k*-th mode of variation is a set of functions indexed by a parameter *α* ∈ ℝ that

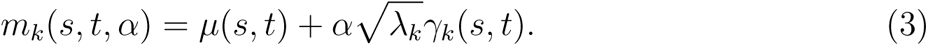

### 3.2 Prediction of Remaining Lifetime

The remaining lifetime prediction (Müller and Zhang 2005) is adjusted to the setup of the function-valued activity process. We denote the lifetime or age-at-death of a subject by *T* and assume that for each subject the daily activity *X* is recorded continuously. We denote the observed activity trajectory for an individual that is still alive at time *T*_0_ by 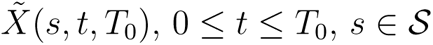, that is

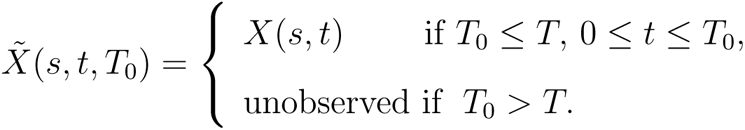

Our goal is to predict the remaining lifetime *T −T*_0_, given 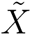(*s, t, T*_0_). Denote the expected remaining lifetime conditional on 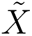 as 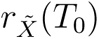, i.e.

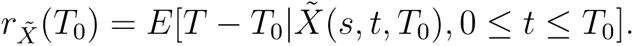

The functional linear regression model (Ramsay and Silverman 2005; Müller and Stadtmüller 2005; Cardot et al. 2003; Müller 2005) is given by

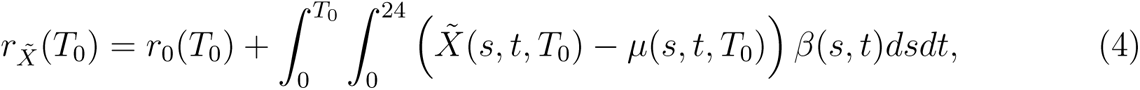

where *r*_0_(*T*_0_) is a nonrandom intercept function, *β*(*s, t*) is the smooth coefficient surface and *µ*(*s, t, T*_0_) is the mean function of 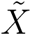(*s, t, T*_0_). The functional linear regression model can be equivalently rewritten as

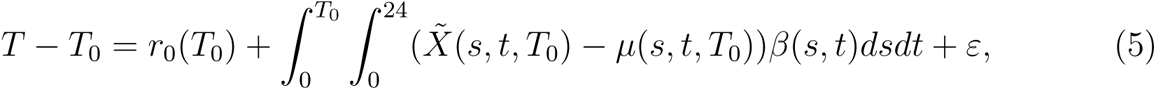

where *ε* is a zero mean finite variance random error. If we represent the coefficient surface *β*(*s, t*) using orthonormal basis *γ_k_* that 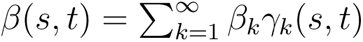, then the model for mean remaining lifetime becomes

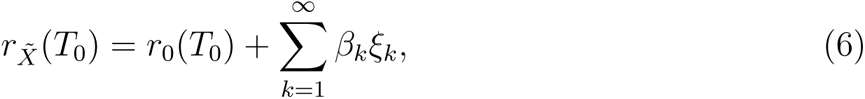

where *ξ_k_*are principal components as in (2).

In order to assess the strength of the association, pseudo *R* square is defined as

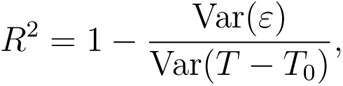

where a larger value of *R*^2^ indicates the predictor explains more of the variability of the response’s remaining lifetime.

### 3.3 Conditional Distributions of Remaining Lifetime

The conditional distribution of remaining lifetime *T − T*_0_ at a current time *T*_0_ of a subject, given an observed trajectory 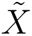 up to time *T*_0_, is defined as 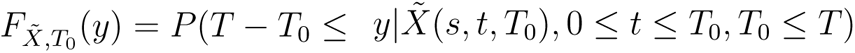. Müller and Zhang (2005) proposed to construct the conditional distribution function of the remaining lifetime by the further assumption that the linear predictor function 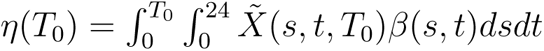 determines the conditional distribution,

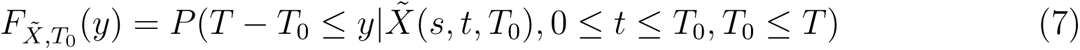

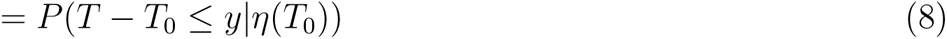

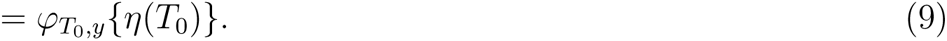

Here the unknown function *φ_T_*_0_*_,y_* (*·*) is assumed to be smooth in *y* and *T*_0_ as their argument. Estimating conditional remaining lifetime distributions then is equivalent to estimating functions *ϕ_T_*_0_*_,y_* {*·*} and *η*(*·*). Nonparametric smooth estimates with boundary kernel (Müller and Wang (1994)) can be utilized to estimate the smooth function *φ*(*·*). See Section 3.3 Müller and Zhang (2005) for more details.

## 4 Results

### 4.1 Early-age Activity Profile and Connection to Diet

The estimated mean process as per (1) of Section 3.1 is shown in Figure 1, along with the mean process under different diet levels. During a 24-hour cycle, the medfly is most active and energetic during periods of illumination as indicated by the light sensor (07:00 - 21:00), and is less active during periods of darkness. With regard to activity dynamics, older medflies exhibit a tendency to rise later. A low nutrition level (C-10) is associated with a relatively delayed activity peak, while a high nutrition level (C-50) is linked to an earlier activity peak.

**Figure 1:**
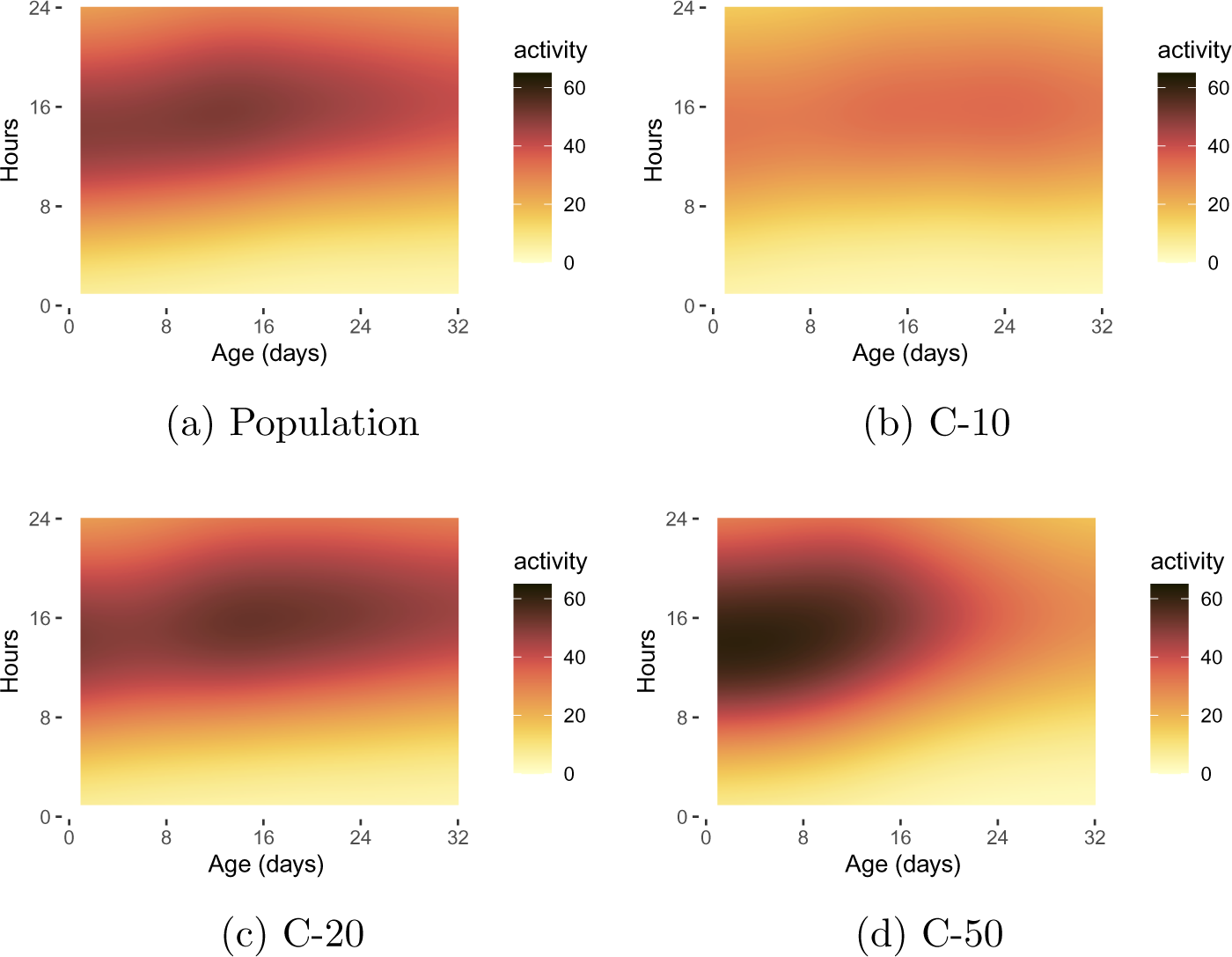
Estimated mean functions of early-age activity, as per (1) of Section 3.1, for population level and three diet levels C-10, C-20 and C-50.

Over 30% of the variation in the early-age activity model (2) is explained by the first three eigenfunctions, as depicted in Figure 11 in the Appendix. The modes of variation of the first three eigenfunctions as per (3) in Section 3.1 are illustrated in Figure 2 to facilitate the interpretation of the eigenfunctions. The predominant variation is reflected in the contrasting patterns of the activity between early and late days and between daytime and nighttime. This indicates that the predominant source of variations are attributed to young medflies during their active daytime hours. The second eigenfunction highlights a contrasting pattern between early and late ages, while the third eigenfunction focuses on the contrast between the activity during day 8 to day 20 (daytime only) and the other periods.

**Figure 2:**
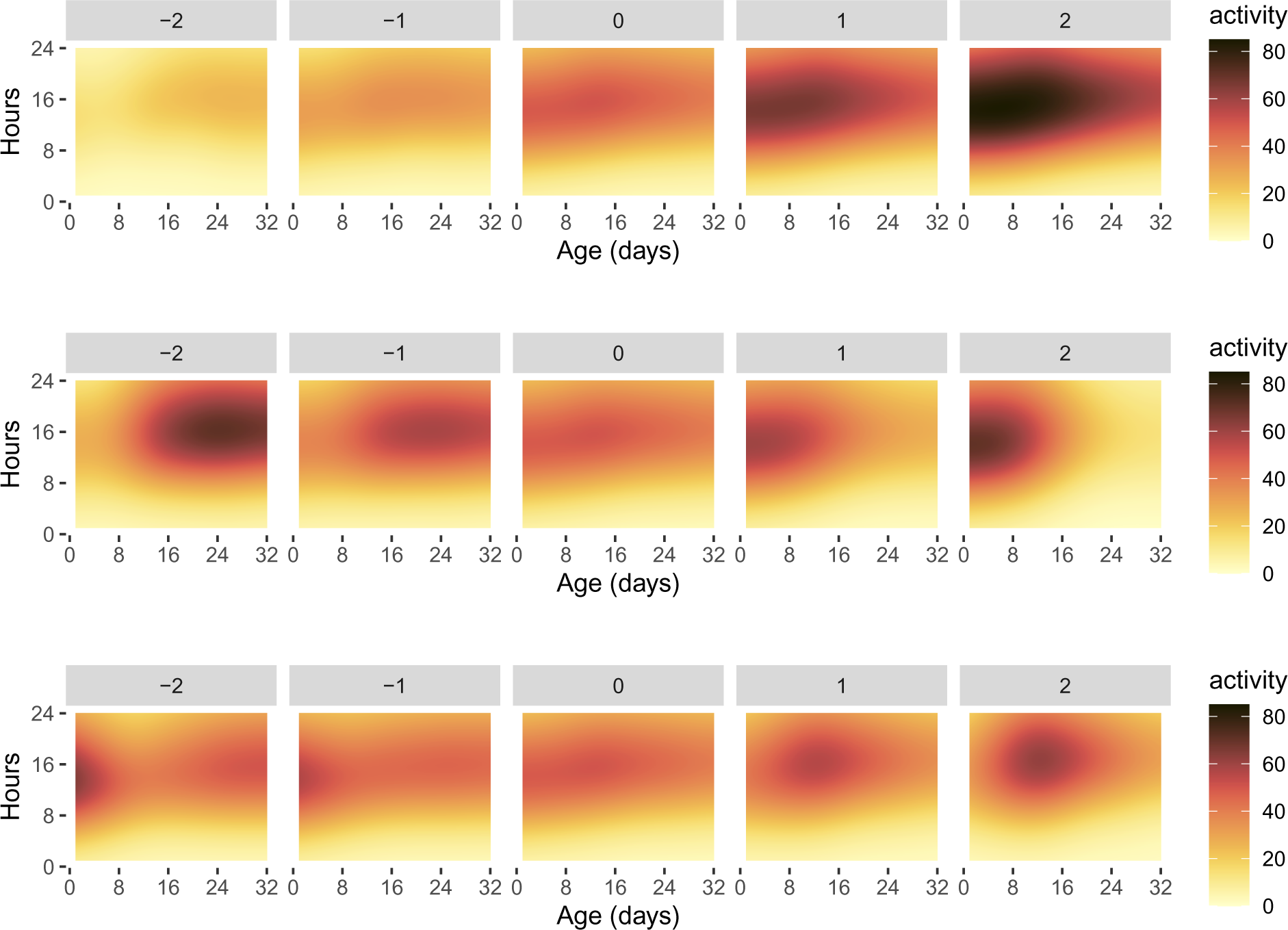
The modes of variation, as per (3) in Section 3.1, vary with the parameter *α* ranging from *−*2 to 2 (left to right). The three rows correspond to the modes of variation of the first three two-dimensional eigenfunctions. The eigenfunctions are illustrated in Figure 11 in the Appendix.

To further investigate the treatment effect on variations of the early-age medfly activity, we fit linear regression models using scores of the FPCA as the response variables and treatment levels as predictors. Table 1 shows the summarised fitted results. To enhance the visualization of the treatment effect, we use pairwise score representations in Figure 10 in the Appendix to facilitate interpretation of the regression models.

**Table 1:** The results of the linear regression using the scores of the FPCA of the early ageage activity as the response variables (top to bottom) and treatment levels as predictors (left to right). The corresponding *p*-values are included in brackets and the *R*^2^ values are listed in the final column. Coefficients with significant *p*-values (*p*-value*<*= 0.05) are shown in bold.

| Response | intercept | C-20 | C-50 | $R^2$ |
| --- | --- | --- | --- | --- |
| $\xi_1$ | **$-243.29(< 0.001)$** | **$295.37(< 0.001)$** | **$389.43(< 0.001)$** | 0.23 |
| $\xi_2$ | **$-132.30(0.01)$** | $50.43(0.455)$ | **$321.96(< 0.001)$** | 0.28 |
| $\xi_3$ | $-50.03(0.24)$ | $27.53(0.63)$ | **$13.31(0.05)$** | 0.06 |

Consistent with the results of the mean activity profiles, a low nutrition level (C-10) is associated with a diminished contrast between daytime and nighttime activity and a relatively late peak in activity. On the other hand, medium nutrition (C-20) and high nutrition (C-50) result in a relatively early peak in activity and an increased contrast between daytime and nighttime activity, with high nutrition (C-50) having a strong connection to high early-age activity.

### 4.2 Relationship between Early-age Activity and Lifespan

In the preceding section, the early-age activity profile of medflies was modeled. This section explores the relationship between the activity profile and longevity and demonstrates that the activity profile can be used as a biomarker for longevity.

We present the results of the functional linear regression model (4) with treatment levels and score representations of the early-age activity profile as predictors. By using the two-dimensional orthonormal eigenfunctions in FPCA, the regression model can be reformulated through equation (6), and the fitted results are summarized in Table 2 where *R*^2^ = 0.21. We also provide the regression model under each treatment level in Table 3. Individuals with medium nutrition (C-20) may expect longer lifespans compared with the other treatments. However, a higher score for the eigenfunction *γ*_1_ (representing high levels of activity in early age and a large contrast between daytime and nighttime activity) can increase the mortality risk for medflies. This effect is particularly pronounced for individuals under a high nutrition level (C-50) as they may have exhausted themselves during their early life or daytime periods. These results indicate that the medfly activity profile can serve as a biomarker of longevity.

**Table 2:** The regression coefficients along with 95% confidence intervals for functional linear regression as per model (6) in Section 3.1 where significant coefficients are bold. Score representations of the early-age activity profile in Section 4.1 together with treatment levels are predictors while the remaining lifespan is the response.

| coefficient | fitted result | confidence interval |
| --- | --- | --- |
| $r_0(T_0)$ | **50.606** | **(40.502,59.811)** |
| $\beta_1$ | **-0.013** | **(-0.034,-0.001)** |
| $\beta_2$ | -0.013 | (-0.030,0.002) |
| $\beta_3$ | -0.005 | (-0.025,0.011) |
| $C_{20}$ | **11.856** | **(0.218, 19.757)** |
| $C_{50}$ | 8.726 | (-6.044,26.473) |

**Table 3:** The regression coefficients under different treatment levels (left to right) as per model (6) in Section 3.2 with score representations of early-age activity profiles in Section 4.1 as predictors (top to bottom) and the remaining lifetime as the response. The *p*-values for each coefficient are also given in the bracket and the significant coefficients (*p*-value *<*= 0.05) are displayed in bold font.

| coefficient | C-10 | C-20 | C-50 |
| --- | --- | --- | --- |
| $r_0(T_0)$ | **49.344**($< 0.01$) | **63.828**($< 0.01$) | **62.975**($< 0.01$) |
| $\beta_1$ | -0.016(0.39) | 0.006(0.52) | **-0.029**(**0.01**) |
| $\beta_2$ | -0.029(0.18) | 0.014(0.40) | -0.008(0.55) |
| $\beta_3$ | 0.017(0.63) | 0.003(0.72) | **-0.028**(**0.05**) |

Figure 3 presents the daily activity curves along with the predicted lifespan in a sample of randomly selected tubes. The results suggest that the prediction of lifespan is in good agreement with the actually observed lifetime. The conditional density curves show the distribution of the predictions, with higher values indicating a higher probability of death at that time.

**Figure 3:**
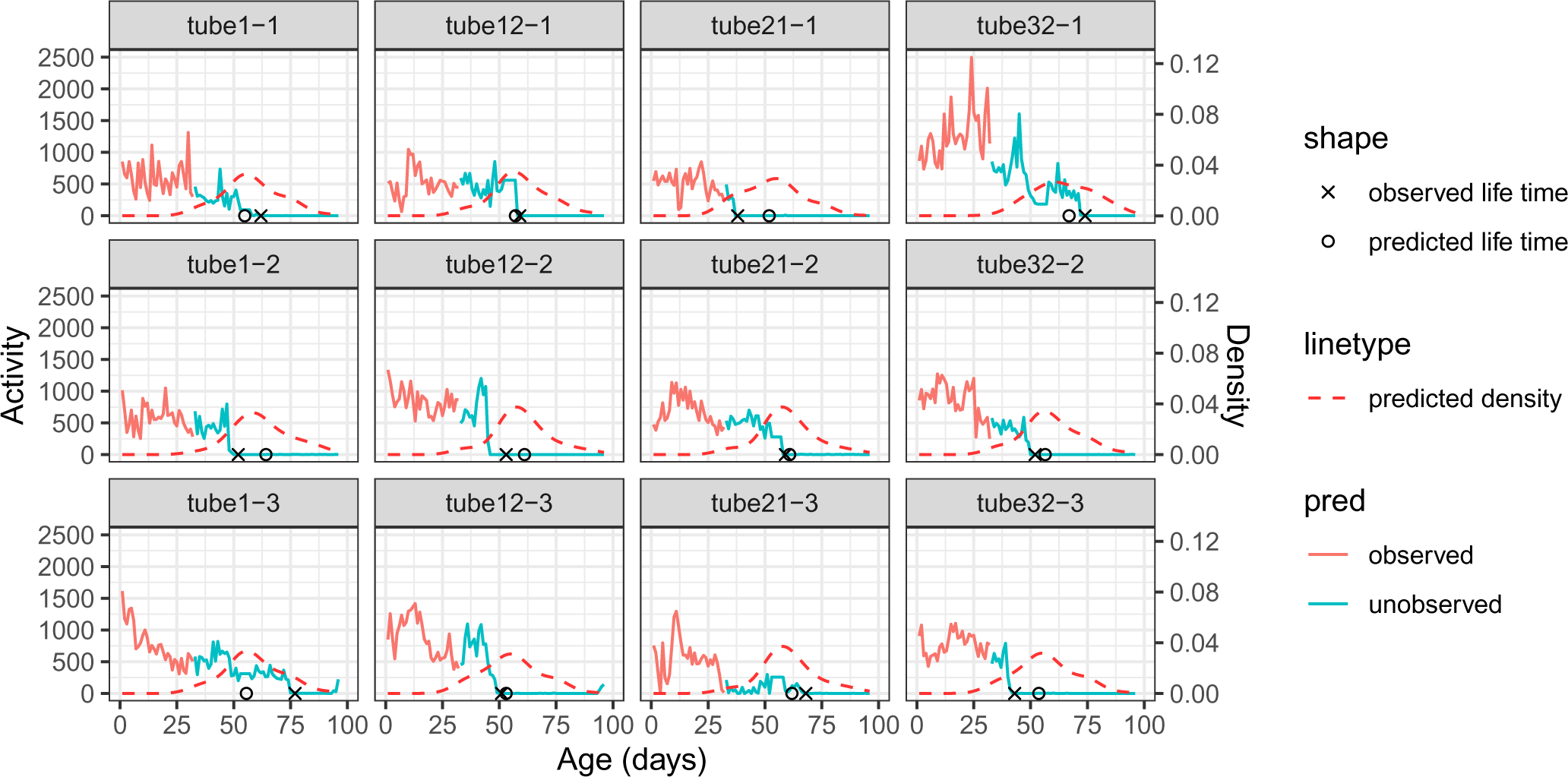
The prediction of the remaining lifespan as per model (6) in Section 3.2, where the score representations of the early-age activity profile in Section 4.1 together with treatment levels are predictors while the remaining lifespan is the response. The figure displays three rows, corresponding to the randomly selected tubes under different treatment levels (C-10, C-20, and C-50, top to bottom). The observed age-at-death is marked as a cross while the predicted age-at-death is marked as a circle for each tube. The predicted conditional density of the remaining lifespan as per model (7) in Section 3.3 is marked with the red dashed line for each tube.

### 4.3 Before-death Activity Profile

In this section, we aim to better understand the activity patterns of medflies before they die. We model the before-death activity as a function-valued stochastic process. The time coordinate is represented by thanatological days (days starting from the day of death), while the repeated measurement coordinate remains unchanged and corresponds to the hour within a day. The mean function and eigenfunctions can be defined and estimated similarly as in Section 3.1

Figure 4(a) shows the population level mean function of the medfly activity modeled in a reversed manner, where the time coordinate starts from the day of death and the hour within the day remains unchanged. The mean function displays the average activity pattern of the medflies, indicating that the medflies tend to be more active during the hours when the light sensor is on (7:00-21:00) and relatively quiet during the hours when the light sensor is off. Additionally, the mean function shows a decline in activity as the medflies approach their death.

**Figure 4:**
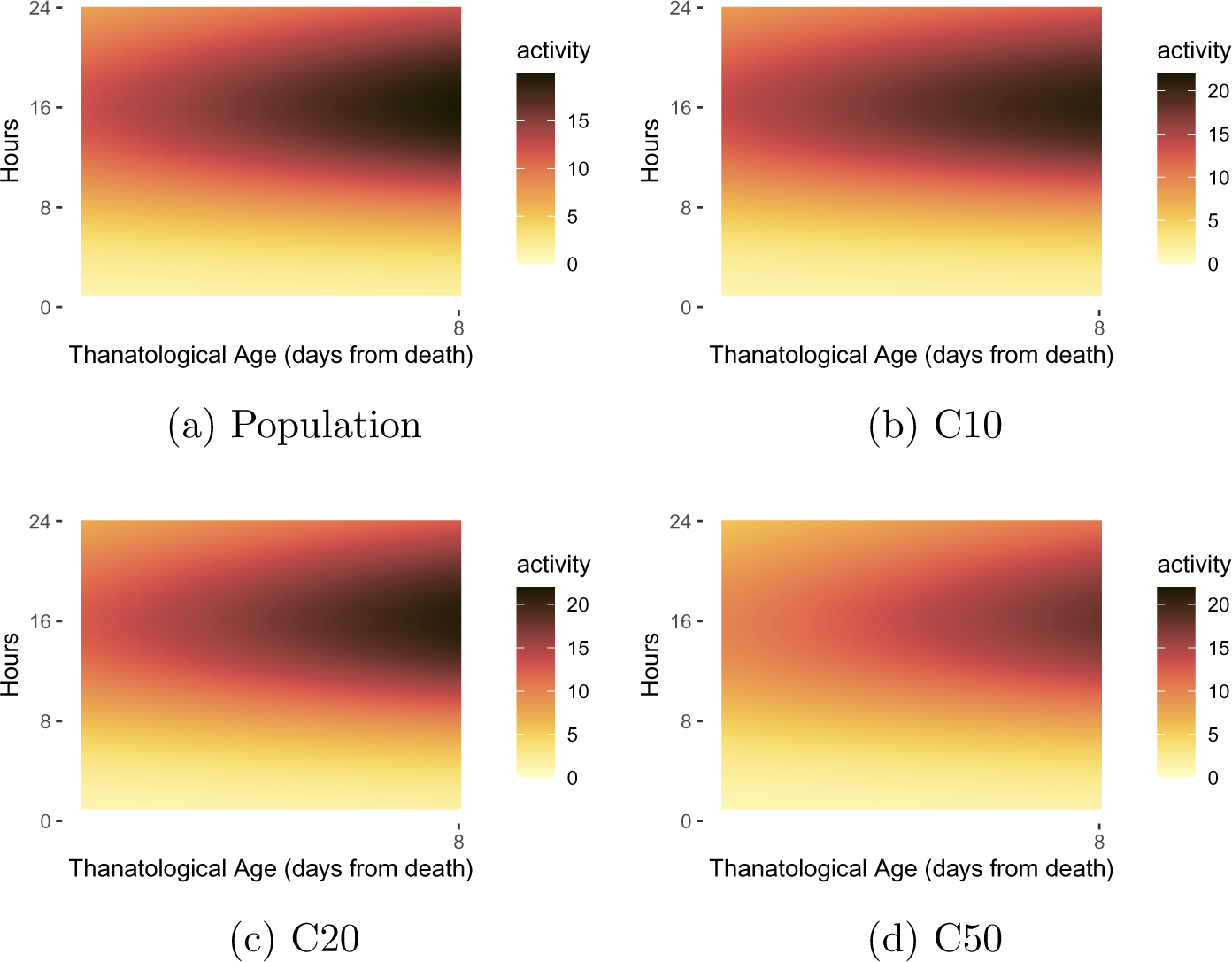
Estimated mean functions of before-death activity profile, as per (1) of Section 3.1, for population level and three diet levels C-10, C-20 and C-50.

Over 50 % of the variation in the before-death activity is explained by the first three two-dimensional eigenfunctions, as depicted in Figure 12 in the Appendix. Modes of variation of the first three eigenfunctions as per (3) in Section 3.1 are illustrated in Figure 5. The primary variation can be seen in the decline of activity during the daytime leading up to death. The second eigenfunction emphasizes the significant level of activity prior to death. The third eigenfunction concentrates on the decrease in activity during the early morning hours.

**Figure 5:**
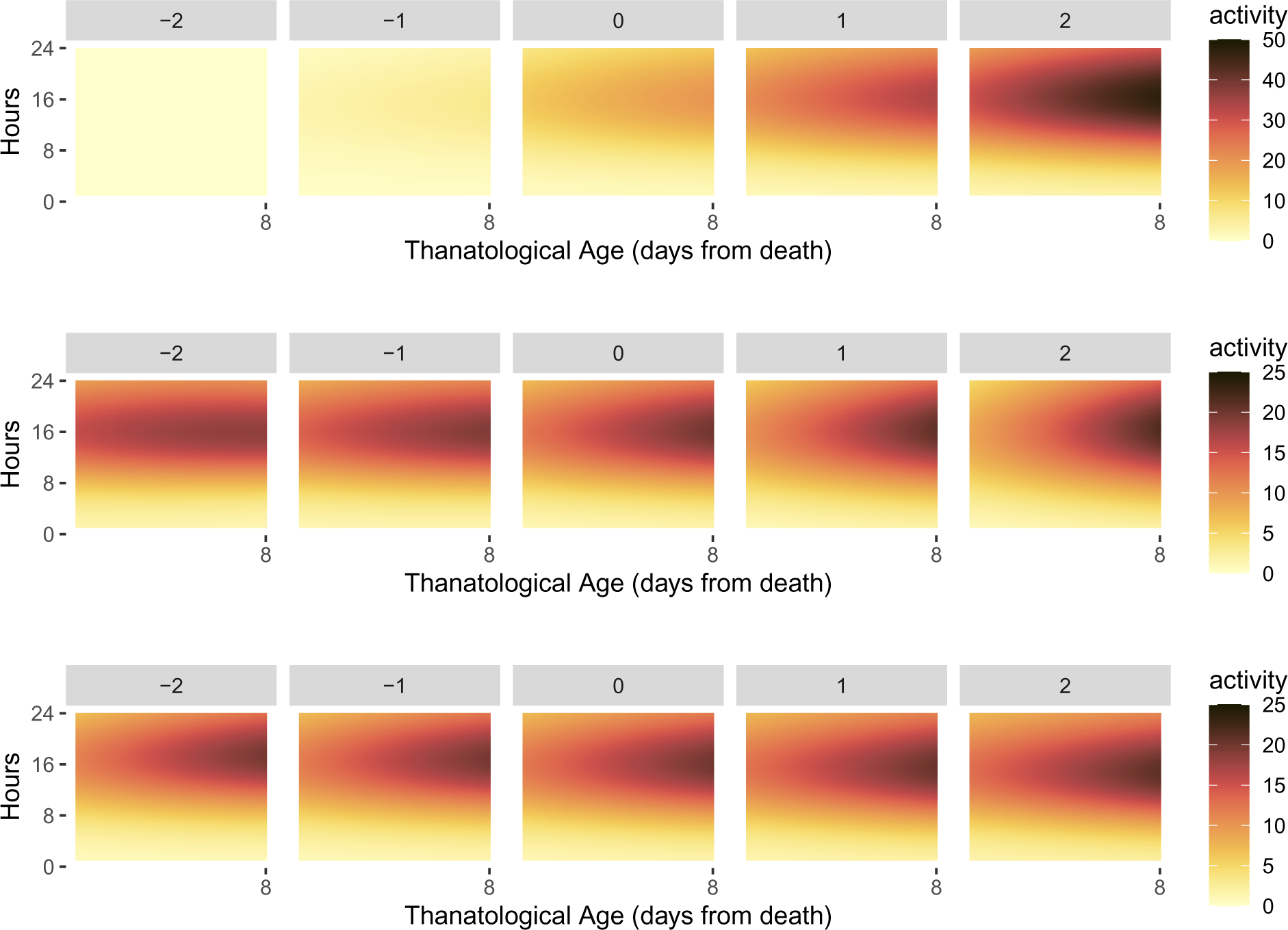
The modes of variation, as per (3) in Section 3.1, vary with the parameter *α* ranging from *−*2 to 2 (left to right). The three rows correspond to the modes of variation of the first three two-dimensional eigenfunctions. The eigenfunctions are illustrated in Figure 12 in the Appendix.

Score representations of the first three eigenfunctions are utilized for downstream cluster analysis. We employ a Gaussian mixture model with a diagonal covariance matrix to identify two underlying clusters, as shown in Figure 6. The first cluster can be connected with the sudden death feature of the medflies, while the second cluster focuses on the dwindling pattern as the medflies approach their death.

**Figure 6:**
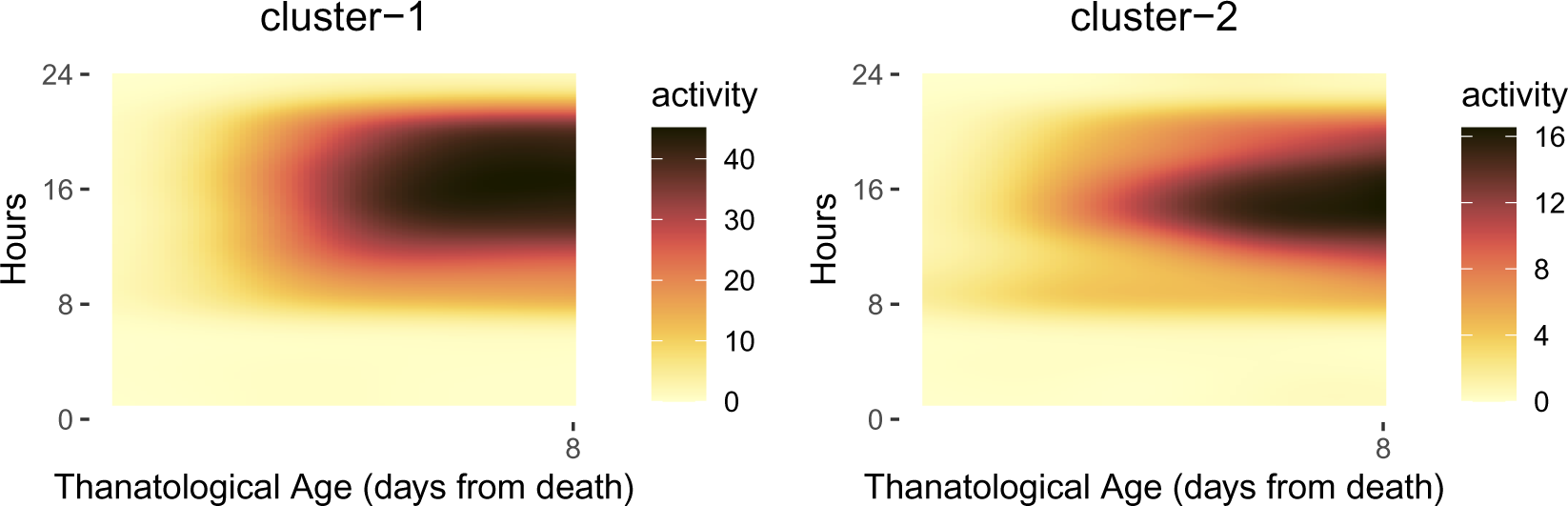
Representatives of the two clusters identified for the activity profiles before death – abrupt end of the activity (left) and dwindling of activity (right).

To gain a deeper understanding of the differences between the two clusters, we randomly select six flies from each cluster and present them in Figure 13 and 14 in the Appendix. For cluster 1, four tubes (24-3, 15-2, 10-1, and 2-2) exhibit a noticeable decline in activity just before death. For cluster 2, all six tubes demonstrate a gradual decrease in activity leading up to death.

## 5 Discussion

We model the daily activity of the Mediterranean fruit fly as a functional-valued stochastic process, taking into account the repeated measurements for each day in the observations. The longitudinal time is represented by the calendar day, and the repeated observations are the recorded times within each day. FPCA is used to effectively demonstrate the variations of the process.

The study identified three distinct patterns of variation in the early-age activity profile and explored their connection to diet. A low nutrition level (C-10) is associated with a diminished contrast between daytime and nighttime activity and a relatively late peak in activity. On the other hand, medium nutrition (C-20) and high nutrition (C-50) result in a relatively early peak in activity and an increased contrast between daytime and nighttime activity, with high nutrition (C-50) having a strong connection to high early-age activity.

Our analysis also provides a strong connection between daily activity in an early age and its subsequent mortality and indicates that the activity profiles can serve as a biomarker. We apply a functional linear regression model that uses the activity within the first 32 days and the treatment received by the medfly to predict its lifespan. Increased mortality risk is linked with higher activity levels and greater contrast between daytime and nighttime activity, particularly in the high nutrition level group (C-50). Individuals receiving medium nutrition (C-20) are predicted to have a longer lifespan compared to those in the low nutrition group (C-10) and high nutrition group (C-50). The predicted lifespan distribution for each subject was also presented, where a higher value of the curve indicates a higher probability of death for the medfly. This shows that the activity profiles can serve as a useful biomarker of longevity.

Additionally, we demonstrate that the technique to model the early-age activity profile can also be applied to the modeling of before-death activity profiles and provides insights into the before-death activity. The use of a Gaussian mixture model cluster analysis identifies two distinct patterns in the before-death activity profiles, allowing for a deeper understanding of the process leading up to death. One cluster is characterized by a slow decline in daily activity and the other is connected with a sudden activity decline before death. Understanding these patterns may help predict future mortality risks and provide valuable information for pest management efforts.

## Acknowledgement

This work is funded in part by a grant from the Center for the Economics and Demography of Aging, UC Berkeley (NIH 2P30AG012839) and in part by an NSF grant DMS-2014626.

## Author Contributions

Han Chen (Methodology, Software, Formal analysis, Writing - original draft). Hans-Georg Müller (Conceptualization, Methodology, Software, Formal analysis, Resources, Writing - original draft, Supervision). Vassilis Rodovitis (Experimental design, Data curation, Writing - original draft). Nikos Papadopoulos (Experimental design, Data curation, Supervision). James R. Carey (Conceptualization, Resources, Writing - original draft, Data curation, Supervision, Funding acquisition)

## Materials & Correspondence

Correspondence and requests for materials should be addressed to Han Chen (email address)

## Data Availability Statement

The data supporting this study’s findings are available from James R. Carey upon reasonable request (email address).

## Appendix

**Figure 7:**
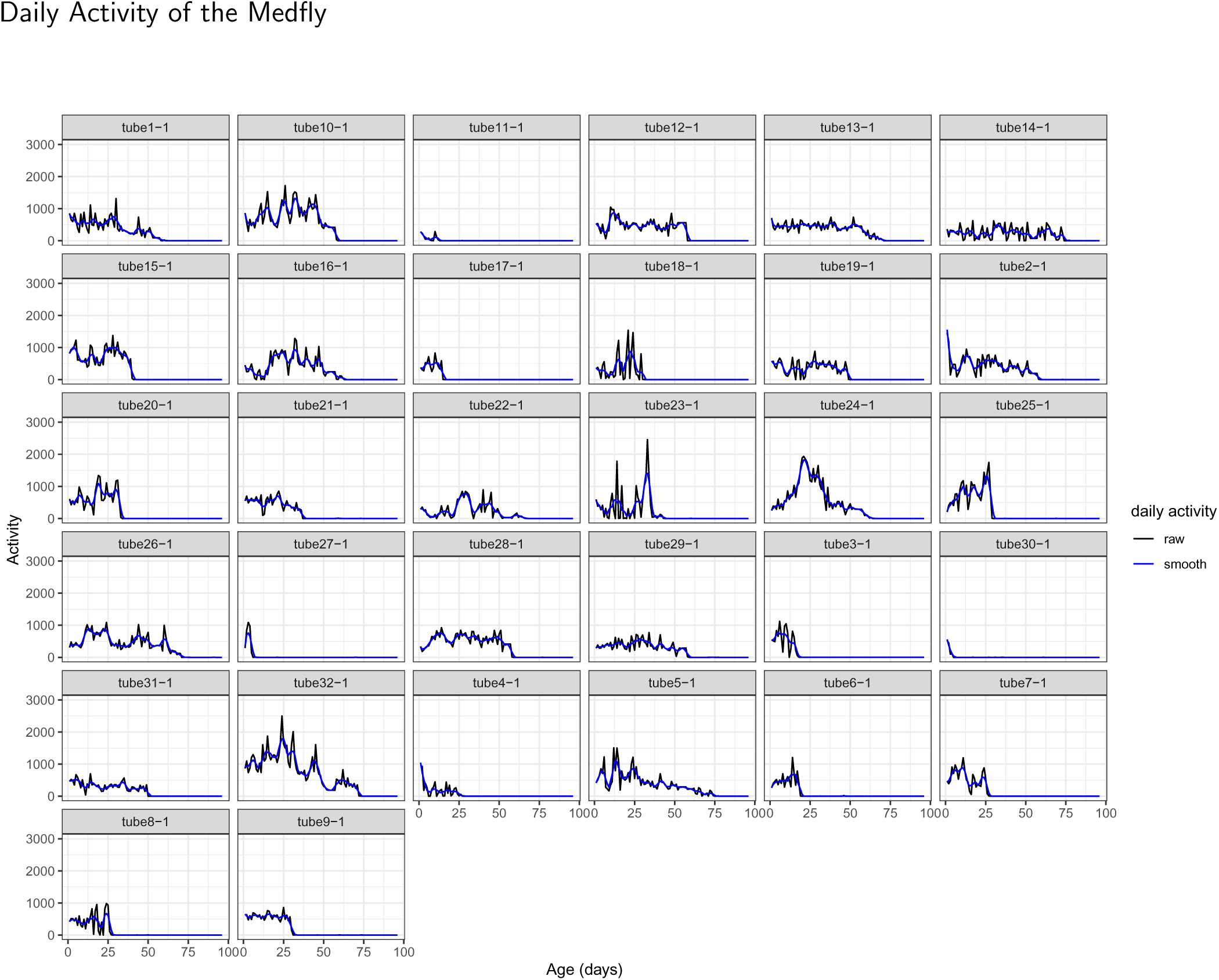
Daily medfly activity under C-10 treatment. The black curves correspond to the raw daily activity trajectory and the blue curves correspond to the local linear smoothing trajectory using Epanechnikov kernel and bandwidth 3.

**Figure 8:**
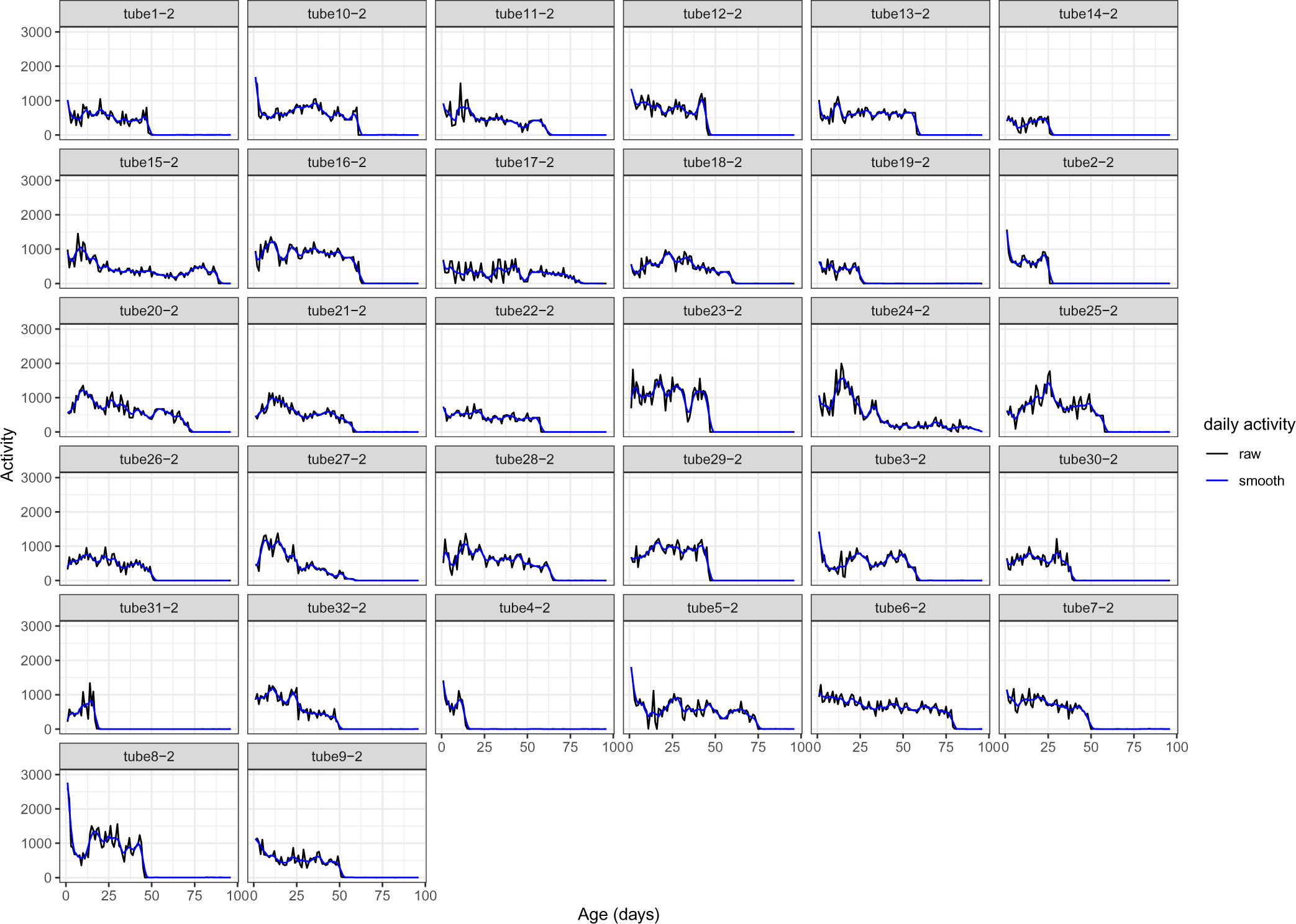
Daily medfly activity under C-20 treatment. The black curves correspond to the raw daily activity trajectory and the blue curves correspond to the local linear smoothing trajectory using Epanechnikov kernel and bandwidth 3.

**Figure 9:**
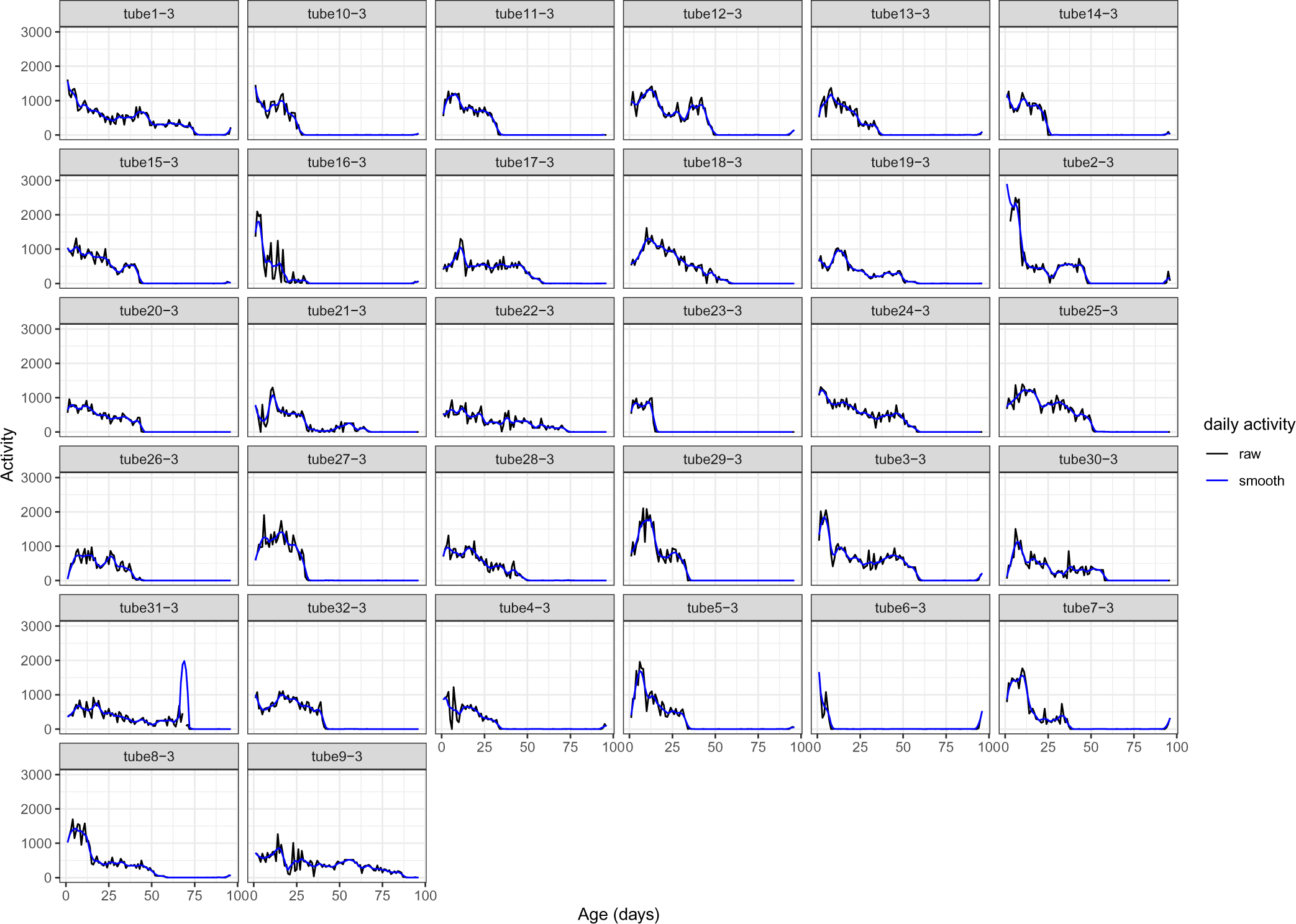
Daily medfly activity under C-50 treatment. The black curves correspond to the raw daily activity trajectory and the blue curves correspond to the local linear smoothing trajectory using Epanechnikov kernel and bandwidth 3.

**Figure 10:**
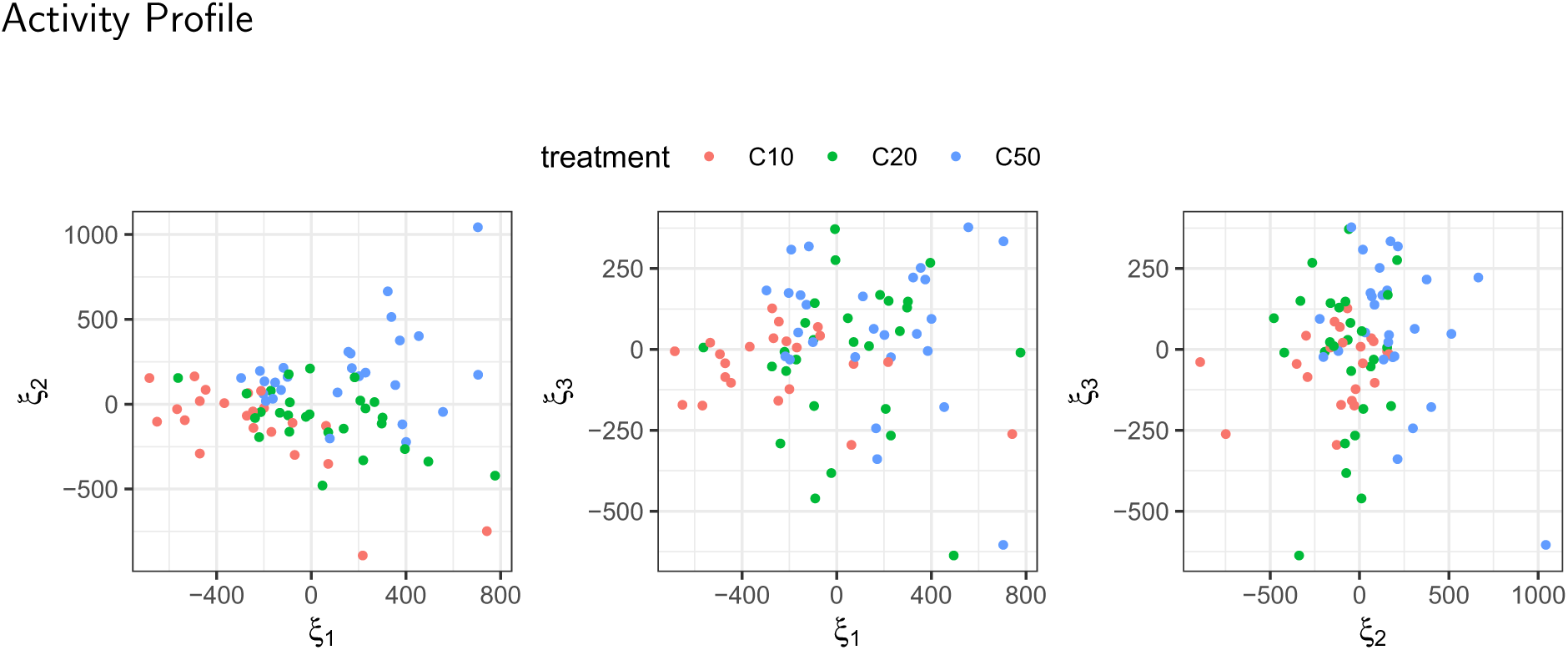
Pair plots of scores of the FPCA of the early age activity in Section 4.1 with the color legend indicates different treatment levels.

**Figure 11:**
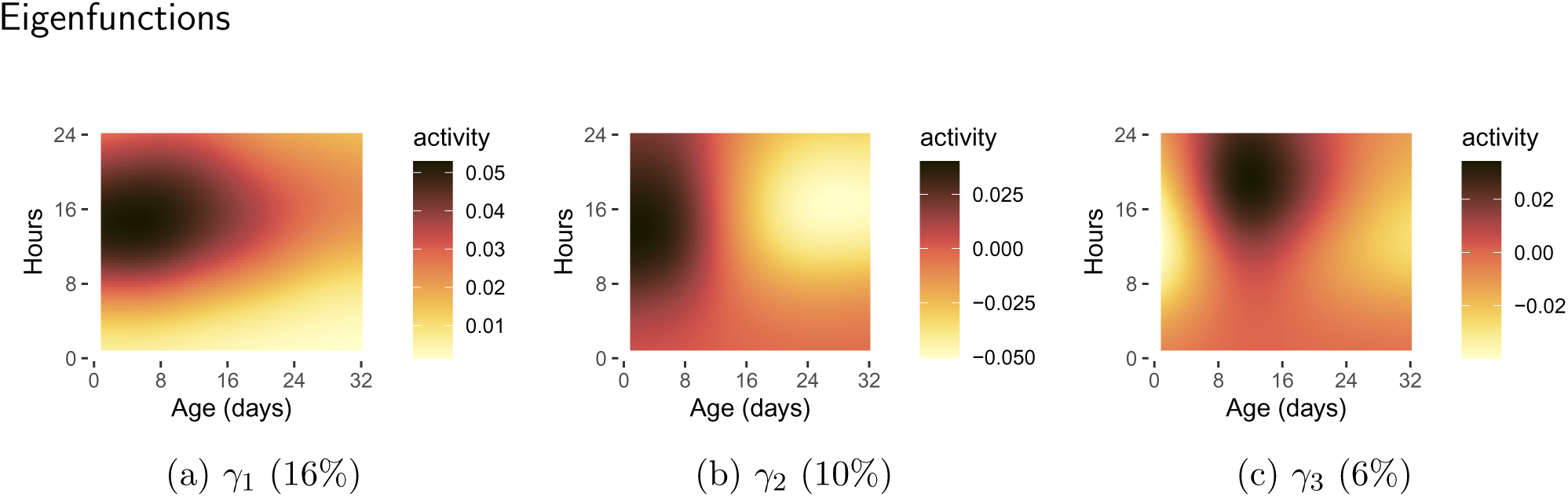
The first three eigenfunctions for FPCA of early age medfly activity as per (2) of Section 3.1. The *Y* -axis represents the coordinate of hours and the *X*-axis represents the coordinate of days from birth. The number in the bracket corresponding to each eigenfunction indicates the fraction of variance explained. Modes of variation for each eigenfunction are illustrated in Figure 2.

**Figure 12:**
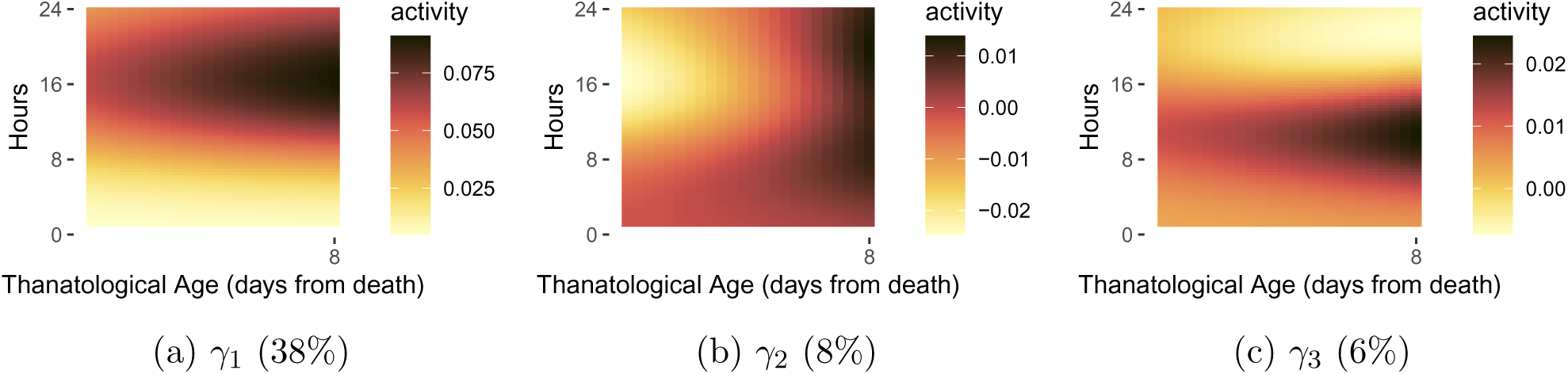
The first three eigenfunctions of FPCA of before-death medfly activity as per (2) of Section 3.1. The *Y* axis represents the coordinate of hours and the *X* -axis represents the coordinate of days from death. The number in the bracket corresponding to each eigenfunction indicates the fraction of variance explained. Modes of variation for each eigenfunction are illustrated in Figure 5.

**Figure 13:**
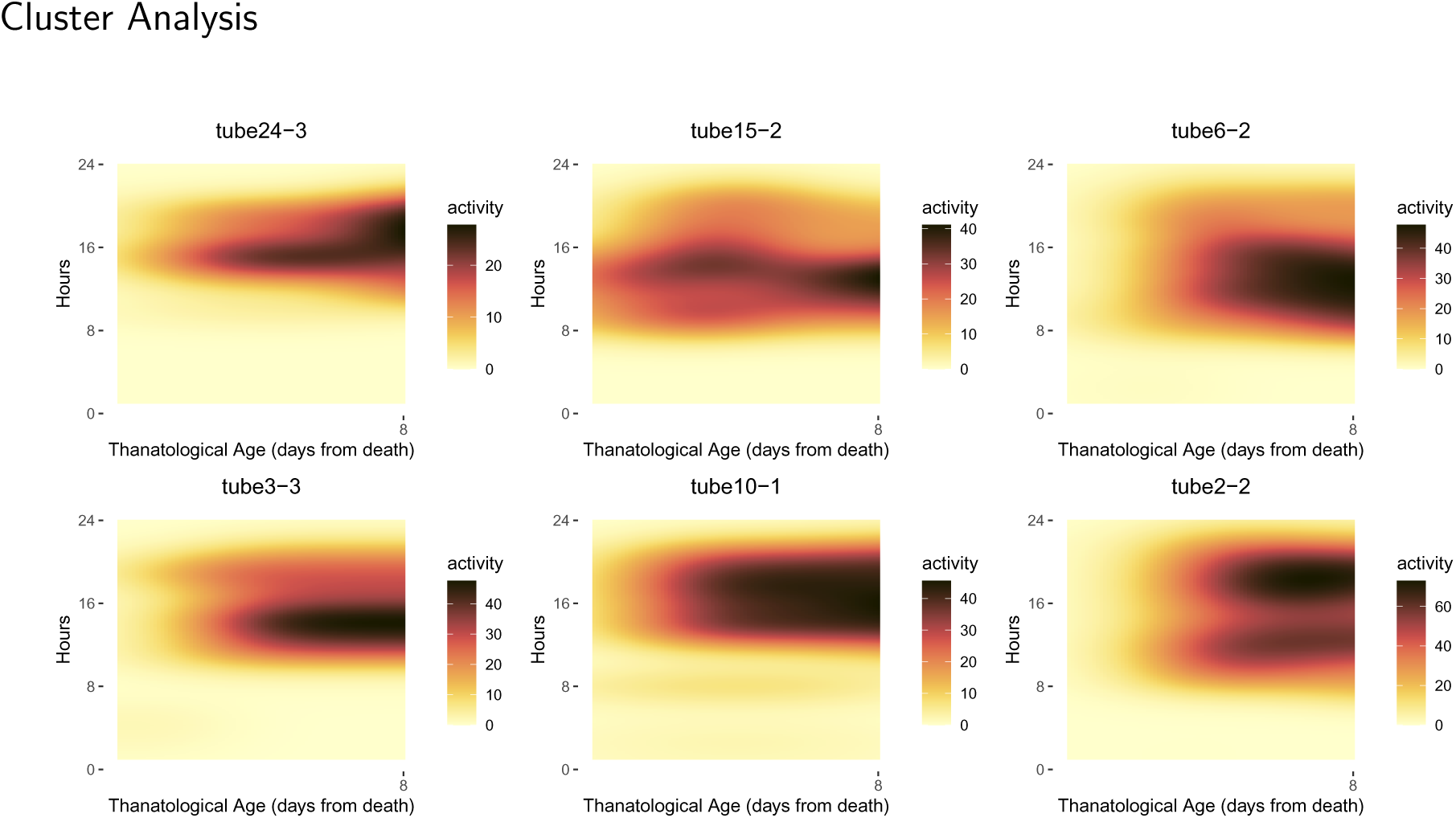
Randomly selected tubes corresponding to cluster 1 as in Section 4.3.

**Figure 14:**
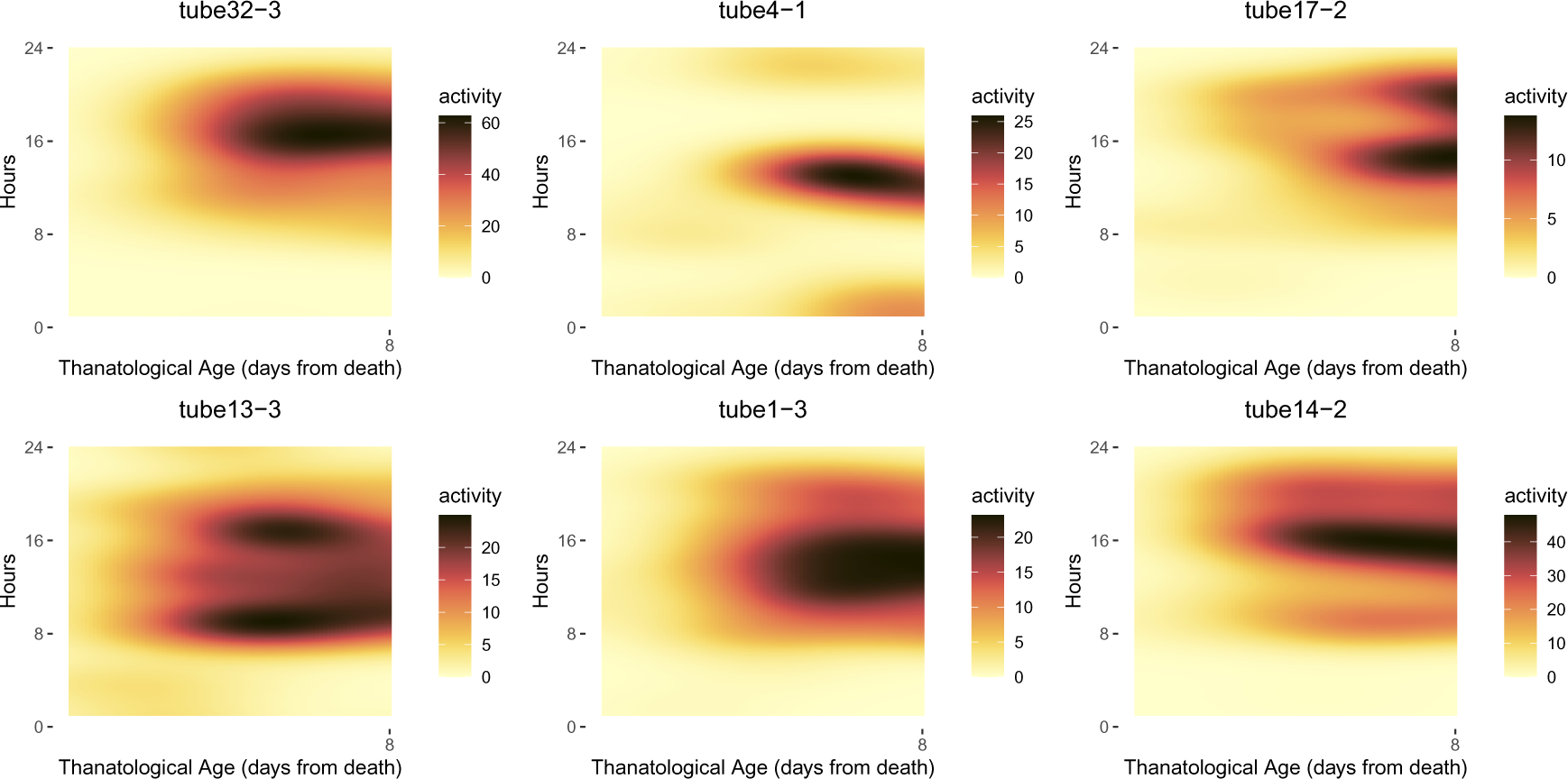
Randomly selected tubes corresponding to cluster 2 in Section 4.3.

